# Imprinting on time-structured acoustic stimuli in ducklings

**DOI:** 10.1101/2021.04.28.441768

**Authors:** Tiago Monteiro, Tom Hart, Alex Kacelnik

## Abstract

Filial imprinting is a dedicated learning process that lacks explicit reinforcement. The phenomenon itself is narrowly heritably canalized, but its content, the representation of the parental object, reflects the circumstances of the newborn. Imprinting has recently been shown to be even more subtle and complex than previously envisaged, since ducklings and chicks are now known to select and represent for later generalization abstract conceptual properties of the objects they perceive as neonates, including movement pattern, heterogeneity, and inter-component relationships of same or different. Here we investigate whether day-old Mallard ducklings (*Anas platyrhynchos*) imprint on the temporal separation between duos of brief acoustic stimuli, and whether they generalize such timing information to novel sound types. Subjects did discriminate temporal structure when imprinted and tested on natural duck calls, but not when using white noise for imprinting or testing. Our data confirm that imprinting includes the establishment of neural representations of both primary percepts and abstract properties of candidate objects, meshing together genetically transmitted prior knowledge with selected perceptual input.

## Background

Under normal circumstances, newborn nidifugous birds, such as ducks and chickens, quickly learn to identify their mother and follow her around for protection, warmth and information on food appearance and location, a phenomenon known as filial imprinting (1).

As a learning mechanism, imprinting is notable because it lacks a specific form of explicit (observable) reinforcement (2) thus exposing both nature and nurture in one sweep, where the mechanism in itself is inherited and narrowly pre-specified, but the content of what is learned is to a large degree left to reflect the circumstances of the newborn. Recent research has shown that, i) sensory pre-dispositions are crucial, facilitating neonates’ orientation towards relevant features of the environment (3,4), while ii) abstract, and logical, relational concepts can be acquired by newly hatched ducklings through imprinting (5). This latter discovery suggests that information learnt during filial imprinting resembles more closely multidimensional vectors that represent potentially abstract concepts than libraries of direct percepts. This interpretation differs from classical models of imprinting, but makes biological sense, considering that successful algorithms for identifying a suitable target object to follow must be robust with respect to scale, perspective, and shape change through body movement (1,6,7). Here we add another dimension of information: the temporal structure of perceived stimuli.

In order to faithfully address behaviour towards an appropriate conspecific, we may expect newborns to form a parental ‘concept’, defined by the three spatial dimensions of stimuli, and by information along the time dimension (in many species, other dimensions, such as an olfactory signature, should be added). Neuroscientists have precisely identified neural codes for spatial information for roughly a century (8), but understanding of brain representations along the fourth dimension, time, while an extremely active area of research (9,10), lags still relatively behind. For this reason, demonstrating that arbitrary temporal features of the environment participate in the representation of imprinting objects would not only enrich understanding of how imprinting works, but also add evidence for how fundamental, widespread and ancestrally old, the ability to keep track of events in time might be.

Drawing inspiration from psychophysical instantiations of timing tasks (11,12), we exposed day-old ducklings to acoustic stimuli that varied in the period of silence between two auditory calls (**Figure 1a**). We used a factorial design with four imprinting treatments resulting from the combination of 2 sound types and 2 temporal intervals of silence, as explained below. Pairs of ducklings were exposed in an arena to 1 of 4 possible stimuli, from an overhanging revolving speaker broadcasting the selected sound stimuli. Each stimulus consisted of two short sounds, or ‘calls’, separated by a silent interval lasting either 0.2 or 1.2 seconds, depending on treatment. Since we know that some environmental features are more salient than others due to innate predispositions (13), the calls themselves were either made from recorded female duck vocalizations (*natural*) or white noise bursts (*artificial*), also dependent on treatment (**Figure 1b**). The property of interest for us was the silent gap, as we were keen to test whether ducklings imprint on the structure of what they hear, as well as on the sounds themselves. Focusing on the duration of a silent gap between two definite percepts controls for the amount of sound energy across stimuli, and exposes sensitivity beyond the sensory features of the stimuli.

**Figure 1.**
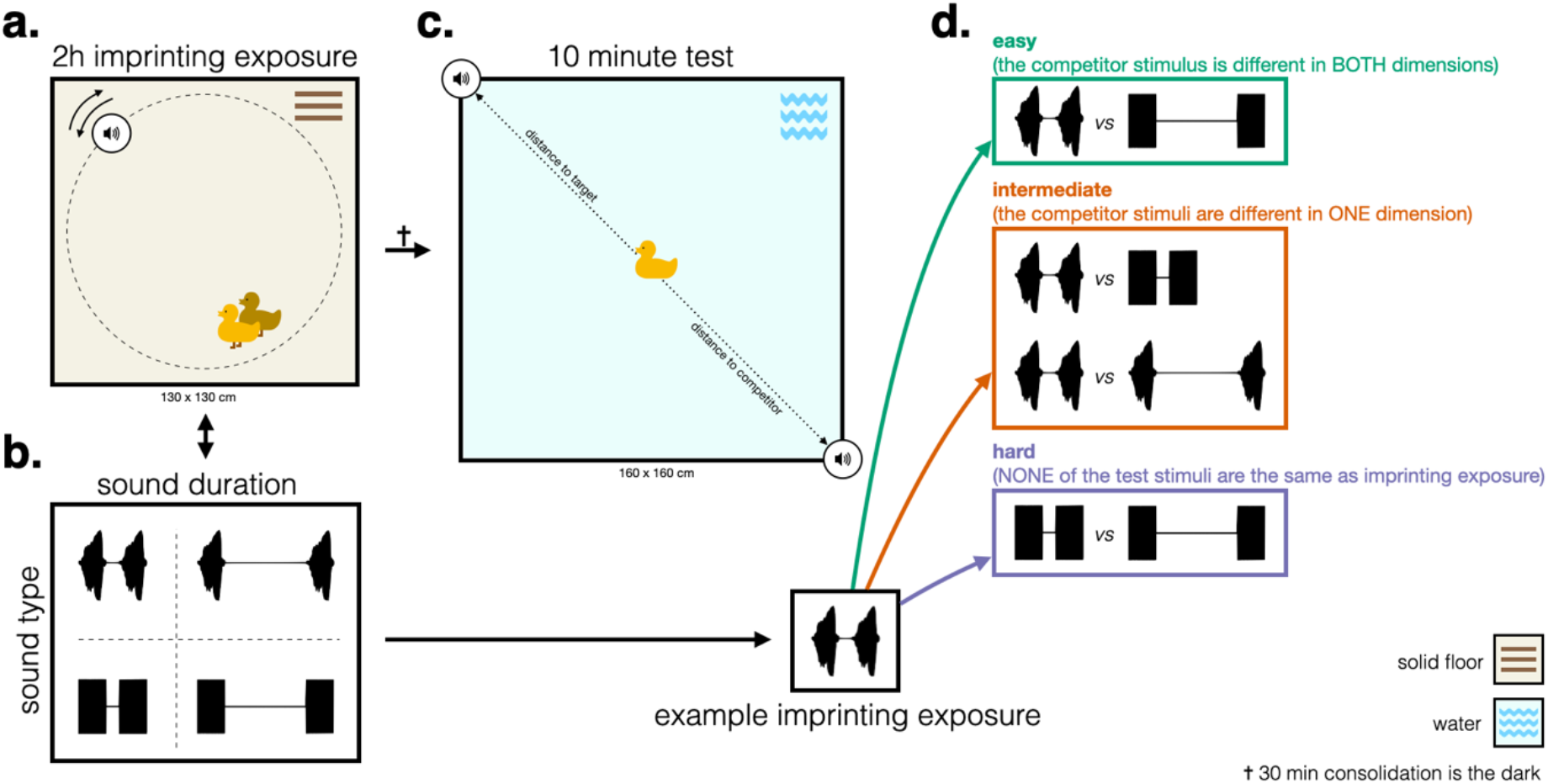
Experimental environment. **a.** Training setup: pairs of ducklings were exposed for 2 hours to a single stimulus, playing from a speaker rotating around the arena **b.** Sound stimuli (normalised amplitude traces) used varied along 2 dimensions, sound time and sound duration. Sounds could be composed of snippets of duck-like call or white noise bursts (rows), separated by either 0.2 or 1.2 seconds silent intervals, for a 1 or 2 seconds total duration (columns) **c.** Testing setup: following a 30-min consolidating period in the dark, single ducklings were tested in a pool with speakers (at fixed locations) playing back two-different stimuli **d.** Example of testing conditions for a particular imprinting exposure stimulus (see Extended figure 1 for a complete depiction of all testing conditions). Colours depict whether the competitor and target test stimuli differed in both (Easy, green) or one (Intermediate, orange) dimensions, or if the target stimulus only retained the same silent gap but not the same sound type as in imprinting exposure (Hard, purple).

## Methods

### Ethics

All procedures were carried out under Oxford University’s animal welfare standards for vertebrates and cephalopods (University of Oxford, 2017) and approved by the Department of Zoology Ethical Review Body. No invasive procedures were carried out.

### Subjects

284 domesticated mallard ducklings (*Anas platyrhynchos domesticus*) of unknown sex were supplied by Foster’s Poultry, Gloucestershire, as eggs, and returned to the supplier as young birds after participating in the experiments. The ducklings were assigned to 4 imprinting exposure groups of varying sizes, that were then split into 16 test conditions (minimum 12, maximum 20 individuals). This allocation was a consequence of varying weekly hatching rates, and ensured that most groups were tested every week.

### Incubation and hatching

Eggs were incubated for 25 days in a Brinsea Ova-Easy 190 incubator in fixed conditions of 37.7 °C and 40% humidity, and moved to a Brinsea Ova-Easy hatcher for the last 3 days of incubation at higher humidity regimes. Hatching took place in the dark to prevent ducklings forming extraneous imprints in the hatching basket. The ducklings remained in the hatching chamber for 12–24h following hatching, as imprinting is most successful during the peak sensitive period between 13 and 40h of age (14).

### Experimental design

Animals were exposed to 1 out of 4 possible acoustic stimuli (**Figure 1b**). Each stimulus could vary in two possible dimensions, sound type (*natural* or *artificial*) and temporal structure (*short* or *long*). In testing conditions (see **Figure 1d** for examples and **Extended figure 1** for full experimental design), animals were tested with a *target* and a *competitor* stimulus, with the *target* stimulus sharing the length of the silent gap with the imprinted one between bursts of either the same or the opposite sound type, and the *competitor* stimulus differing from the target in either two dimensions (sound type and gap duration) or only one of them. Test conditions thus comprised 3 levels of difficulty, categorised as ‘easy’ (competitor different in both dimensions), ‘intermediate’ (competitor different in one of the dimensions) and ‘hard’ (both stimuli composed of a sound type not used in imprinting exposure). In the hard test condition the *target* stimulus shared only the temporal profile with the stimulus to which animals had been exposed during the imprinting phase.

### Stimulus design

Sound stimuli were either composed of natural female duck calls or white noise bursts (0.4 seconds). Each stimulus included two sounds of the same type separated by 0.2 (short stimuli) or 1.2 (long stimuli) seconds silent intervals. Overall stimuli had either 1 (short stimuli) or 2 seconds (long stimuli) duration. All 4 individual sound stimuli (see Experimental design and **Figure 1b**) were generated using Audacity *software* (audacityteam.org). The female adult mallard quack sound burst was extracted from the Macaulay Library at the Cornell Lab of Ornithology audio file (ML 133222). Session long sequences of stimuli were compiled into mp3 files using custom Matlab code (2020a, Mathworks). Stimuli were separated by variable inter-stimulus silent intervals drawn from a normal distribution (u=15, σ=5 seconds).

### Imprinting exposure

Following previous experiments from our laboratory (5,15), we combined priming, shown to produce improved imprinting responses in chickens and ducklings (16,17) with 2h long imprinting exposure periods. Pairs of ducklings were exposed in an arena (130 x 130m, **Figure 1a**) to an overhanging wireless speaker (EasyAcc, model LX-839) playing back one of 4 possible stimuli (see Experimental design, above). The speaker was suspended 15 cm from the floor by an invisible fishing line attached to a rotating boom. Each revolution lasted approximately 40 s, with a diameter of 1 m, as movement has been shown to enhance imprinting (18). There were 4 possible imprinting exposure conditions: natural short, natural long, artificial short and artificial long stimuli (**Figure 1b**). Following imprinting exposure, duckling pairs were placed in a dark chamber for a 30-min retention interval.

### Testing

Individual ducklings were tested for 10 minutes in a pool (180 x 180 m, **Figure 1c**), with stimuli placed in 2 diametrically opposed fixed locations (position of target and competitor balanced across subjects). There were 16 testing conditions (see **Figure 1d** and **Video 1** for an example, and **Extended figure 1** for the complete experimental design).

### Data acquisition, processing and analyses

Video was recorded using Sony wireless 4K action cameras (FDR-X3000 R) at 30 Hz. A colour thresholding method was implemented using custom *Bonsai* code (19) to track the position of the duckling and each speaker. Position data was downsampled to 1Hz, rotated and normalised relative to the position of the sound speakers using custom Matlab code (2020a, Mathworks), so that speakers were always at the x,y positions [-1, 0] and [1, 0] (see **Extended figure 2** for individual examples). The first 20 seconds of each test were discarded to allow the duckling to get used to being in water for the first time. We then computed the second-by-second Euclidean distance between the duckling and each of the speakers (see dashed lines in **Figure 1b** for schematics). For every animal we computed a preference index ‘Delta’ by subtracting the average Euclidean difference for the competitor speaker from the average Euclidean distance to the target one, so that positive deltas correspond to shorter average distances to the target and negative ones indicate a preference against it. Violin plots included in Extended figure 3 were created using (20).

A 3-way ANOVA with imprinting exposure sound-type (2 levels: natural and artificial), silent gap duration (2 levels: short and long), test difficulty (4 levels: easy, two intermediate; hard) and all pairwise interactions between these factors was used to test for preference (R aov function):

~~~
Preference ~ SoundType + SilentGapDuration + TestDifficulty + [all pairwise interactions]
~~~

The interaction plot shown in Extended figure 4 was built using R function

~~~
(cat_plot).
~~~

### Data and code accessibility

All data can be found as electronic supplementary material.

The code that supports the findings of this study is available at https://github.com/PTMonteiro/MonteiroHartKacelnik_2021.git

## Results

Ducklings (n = 284) were exposed (imprinting exposure) to a target stimulus on solid ground and tested in a water pool (**Figure 1c**), to mimic common natural circumstances for imprinting and locomotory following mode. During preference tests the speakers broadcasted the sound stimuli from 2 diametrically opposed fixed locations suspended a few centimetres above opposite corners of the pool. In all treatments one stimulus (*the target*) had the same silent gap between two short sounds as the one used during imprinting exposure, but the gap occurred between either the same or a novel sound type. The other speaker emitted the *competitor* stimulus, that could differ from the imprinting stimulus in either both dimensions (sound type and gap duration) or only one of them (see **Figure 1d** for examples and **Extended figure 1** for full experimental design), thus exploring different levels of generalization.

The factorial combination of 2 sounds and 2 temporal gaps resulted in 4 imprinting exposure groups, each of them divided in 4 testing groups, thus generating 16 testing subgroups (**Extended figure 1**), and allowing for the discrimination of the differential effects of imprinting to a time interval and inherited preferences between natural and artificial sounds. The 16 testing subgroups posed 3 levels of difficulty, depending on how distant they were from the original imprinting exposure conditions. In the 4 ‘Easy’ subgroups ducklings’ faced a *target* stimulus equal to the corresponding imprinting stimulus, and a *competitor* stimulus different to the *target* in both silent gap duration and sound type. In the 8 ‘Intermediate’ subgroups the *target* was still the same as the imprinting stimulus, but it was tested against a *competitor* different in either the gap duration or the sound type, but not in both. Finally, in the 4 ‘Hard’ subgroups the imprinting stimulus was not present at the time of testing: both the *target* stimulus and the competitor were made of novel sounds, but in the *target* the duration of the silent time gap was as during imprinting exposure, while in the *competitor* it was different, so that only temporal structure (duration of gap between the ‘calls’) served as discriminant. This last condition was the most critical, requiring ducklings to generalize from a silent temporal gap between 2 sounds heard at the time of imprinting to the same temporal gap between novel sounds at the time of testing. This condition resembles the abstract relational concept imprinting tested previously with visual stimuli (5).

**Figure 2** (see **Extended figure 3** for individual animal data) shows that ducklings preferred the target stimulus when the target stimulus was constructed with natural duck calls (white bars). However, while this shows some sensitivity to the temporal relation, the kind of sound seems to play a major role. In the ‘Easy’ conditions, ducklings preferred the stimulus composed of natural sounds, regardless of the gap length, as shown by positive white and negative black bars, respectively (**Figure 2**, green panel). For ‘Intermediate’ difficulty (**Figure 2**, orange panels), the *target* stimulus had the same sound and gap as that during the imprinting exposure, and the *competitor* differed in either sound or gap duration, but not in both. Here, when the test stimuli differed in sound but not gap, the ducklings preferred the natural sound regardless of their exposure treatment, and when the test stimuli differed in the temporal gap but not in the sounds, ducklings preferred the target when composed of natural sounds, but were indifferent when composed of artificial sounds. On their own, the results of the intermediate difficulty would be consistent with an ability to imprint on the temporal structure of natural but not artificial sounds, perhaps to be expected if natural sounds are more attentionally salient. In the ‘Hard’ condition (**Figure 2**, purple panel), both test stimuli were composed of a novel sound, but only the *target* had the temporal gap experienced during imprinting exposure. The interaction between *sound type* used during imprinting exposure and *difficulty* was highly significant, showing that increasing *difficulty* lessened the effect of *sound type* (F_3,271_=8.049, P<0.001; Figure 2). The interaction plot in **Extended figure 4** shows that the interaction is driven by the preference for targets composed of natural sounds decreasing with *difficulty*, while *difficulty* had no definite effect on preference for targets composed of artificial sounds.

**Figure 2.**
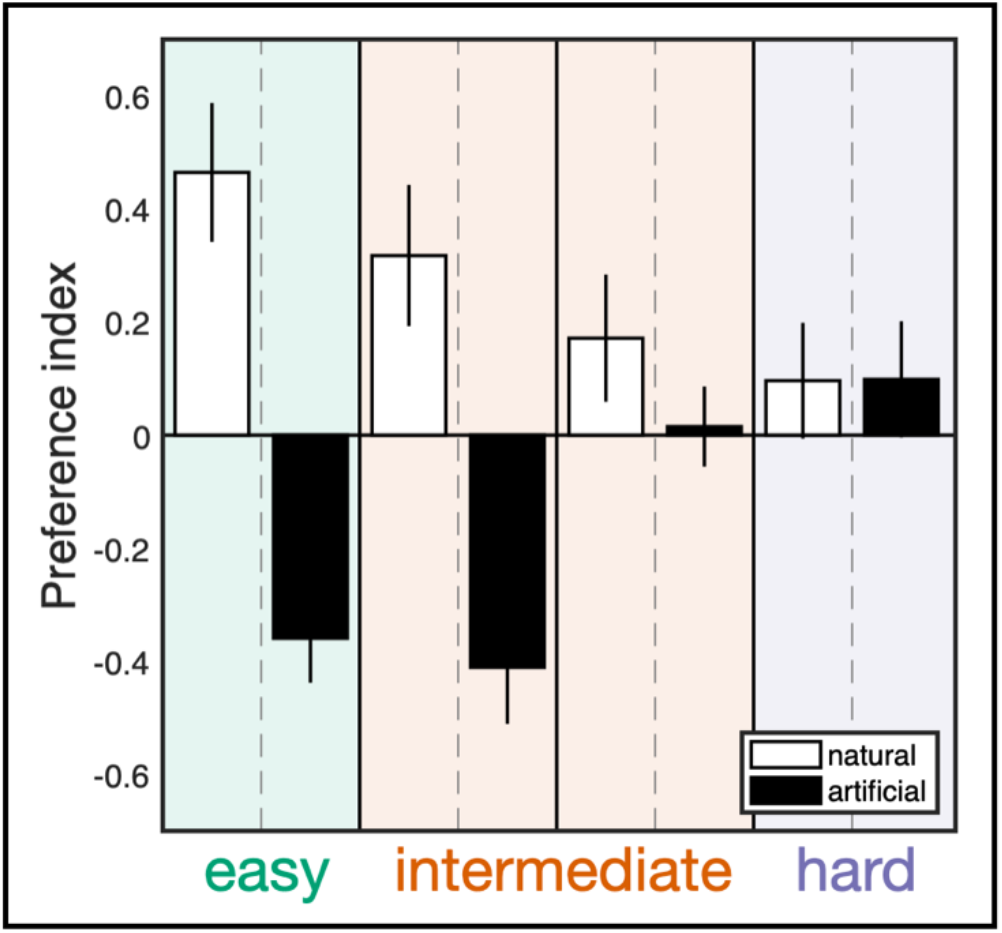
Preference in tests. Preference index (means ± sem; n = 284, see main text for details). Same colour scheme as in Figure 1. White and black bars distinguish imprinting exposure conditions where natural or artificial sounds were used, respectively.

## Discussion and Conclusions

Overall our results point to the complexity of information processing during imprinting, even if they do not reliably establish whether imprinted ducks generalize the temporal intervals between acoustic stimuli across arbitrary sound types. When imprinted on natural duck calls or white noise, ducklings reliably prefer to follow natural calls regardless of their temporal organization. However, when animals face a choice between natural calls with varying temporal structure, they do prefer the temporal structure they have experienced in the imprinting phase. They did not display the same sensitivity to time separation of calls when their imprinting exposure was with acoustic stimuli made out of white noise.

In summary, the existence of sensitivity to an organizational property of stimuli, together with a strong effect of prior predispositions, confirms the view that even in this specialized, unreinforced form of learning the representation of information in the brain takes the form of concept vectors with multiple attributes that make it possible generalization towards stimuli that are identified by abstract rather than direct perceptual properties. With hindsight, and by contrasting our methods with protocols more widely used with research on imprinting in chickens (21–26), it is to be expected that clearer evidence of generalization would result from longer imprinting exposures. We only offered ducklings a period of about 2 hours of imprinting before testing for preference, while most studies in chicken use exposures well over a full day. It is probable that our subjects were still in the midst of their sensitive period and under those circumstances they may even have some level of preference for ‘novel’ stimuli, so that even if they did discriminate the temporal structure of the two choice signals, our preference index would not have detected because they would ‘explore’ the competitor (7). An increase in experimental power could be achieved without increasing the total number of subjects, by taking ducklings’ inherited preference for the natural calls as proven, and testing participants only for sensitivity to temporal structure. Additionally, the duration of imprinting exposure periods would probably strengthen the preferences, but would necessarily increase the involvement time of each subject and thus reduce efficiency.

Our finding is consistent with, but does not prove, the hypothesis that conceptual imprinting extends to the temporal relation between acoustic stimuli, further broadening the sensory domains of this intriguing learning phenomenon.

## Supporting information

Video1.Example animal trajectory with overlaid automatic tracking.Corresponds to top left panel in Extended Figure 2.

## Authors’ contributions

T.M. and A.K. designed the experiment; T.M. ran the experiment; T.M. and T.H. analysed the data; T.M. and A.K. wrote the first draft of the paper; all authors revised the manuscript, approved the final version of the manuscript, and agree to be held accountable for the content therein.

## Acknowledgements

We thank Antone Martinho-Truswell for technical advice, Eli Versace, Dora Biro, Pedro Lacerda, Filipe Rodrigues, Margarida Pexirra, Andreia Madalena and Miguel Santos for discussions of earlier versions of this work, the Wytham Field Station staff for logistical support, Mary Pocock from Fosters Poultry for supplying our eggs and collecting our ducklings back to the farm, and The Macaulay Library at the Cornell Lab of Ornithology for providing access to the audio recording used to design our natural experimental stimuli. AK is grateful for the support of the Deutsche Forschungsgemeinschaft (DFG, German Research Foundation) under Germany’s Excellence Strategy – EXC 2002/1 “Science of Intelligence” – project number 390523135.

## Supporting material

**Extended figure 1.**
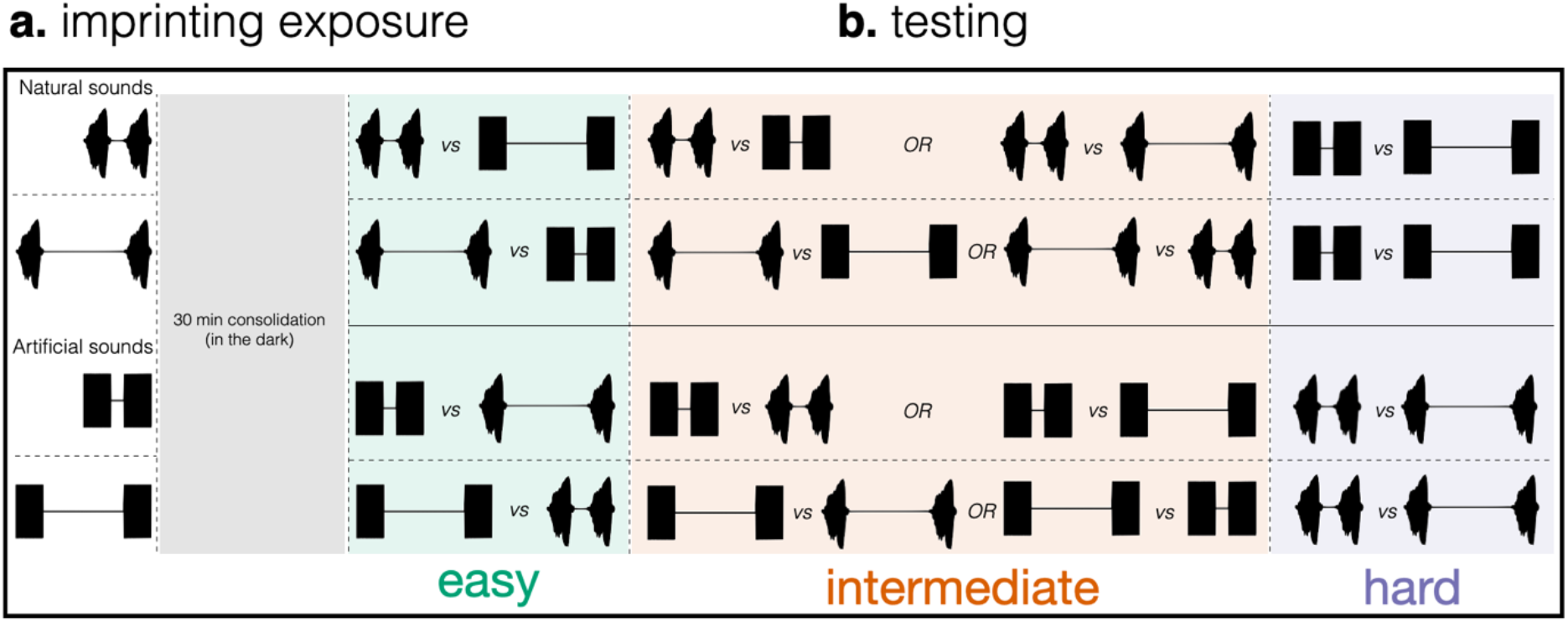
Full experimental design. **a.** Imprinting exposure (training) conditions. Animals were exposed to one of 4 possible stimuli (normalised amplitude traces). **b.** Testing conditions. Following a 30-min consolidation period in the dark (grey shaded rectangle), each animal was then exposed to one test condition, out of 4 possibilities, composed by a target and a competitor stimulus. Across test difficulties the competitor was different from the preexposed stimulus in both sound and length of the gap (Easy), in only one of them (Intermediate), or, for the most difficult condition (Hard), both stimuli were composed of a sound type different from the pre-exposed stimulus, but one of them (the target) shared the same temporal gap (see Figure 1d for an example). Colour scheme as in Figure 1.

**Extended figure 2.**
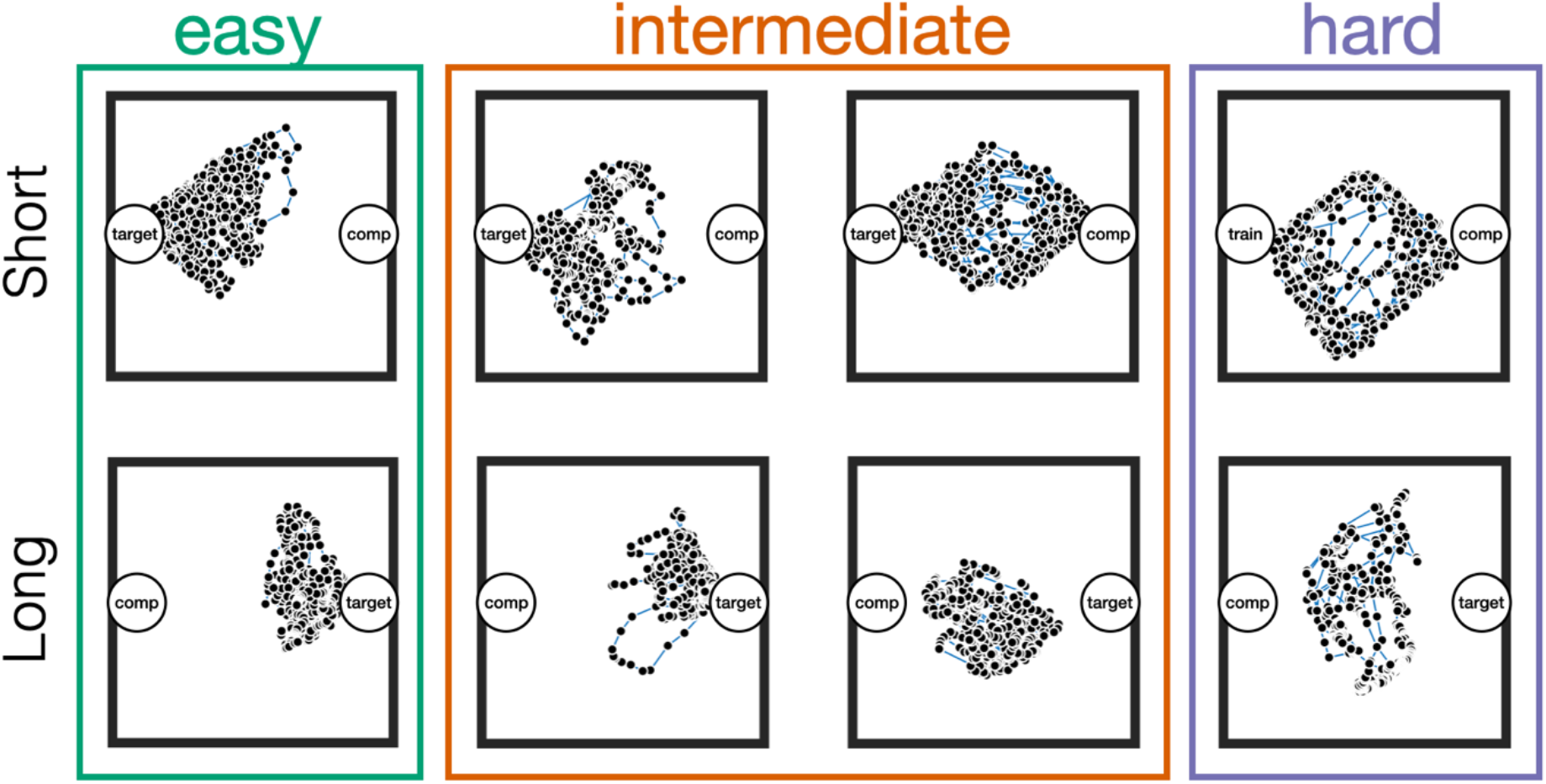
Examples of individual animal trajectories in tests where animals had imprinting exposure to natural sounds with different silent gap parameters (top row, short; bottom row, long). Each black dot shows the position of the animal at 1Hz resolution, with blue lines connecting adjacent positions. Coloured rectangles use the same colour scheme used in previous figures.

**Extended figure 3.**
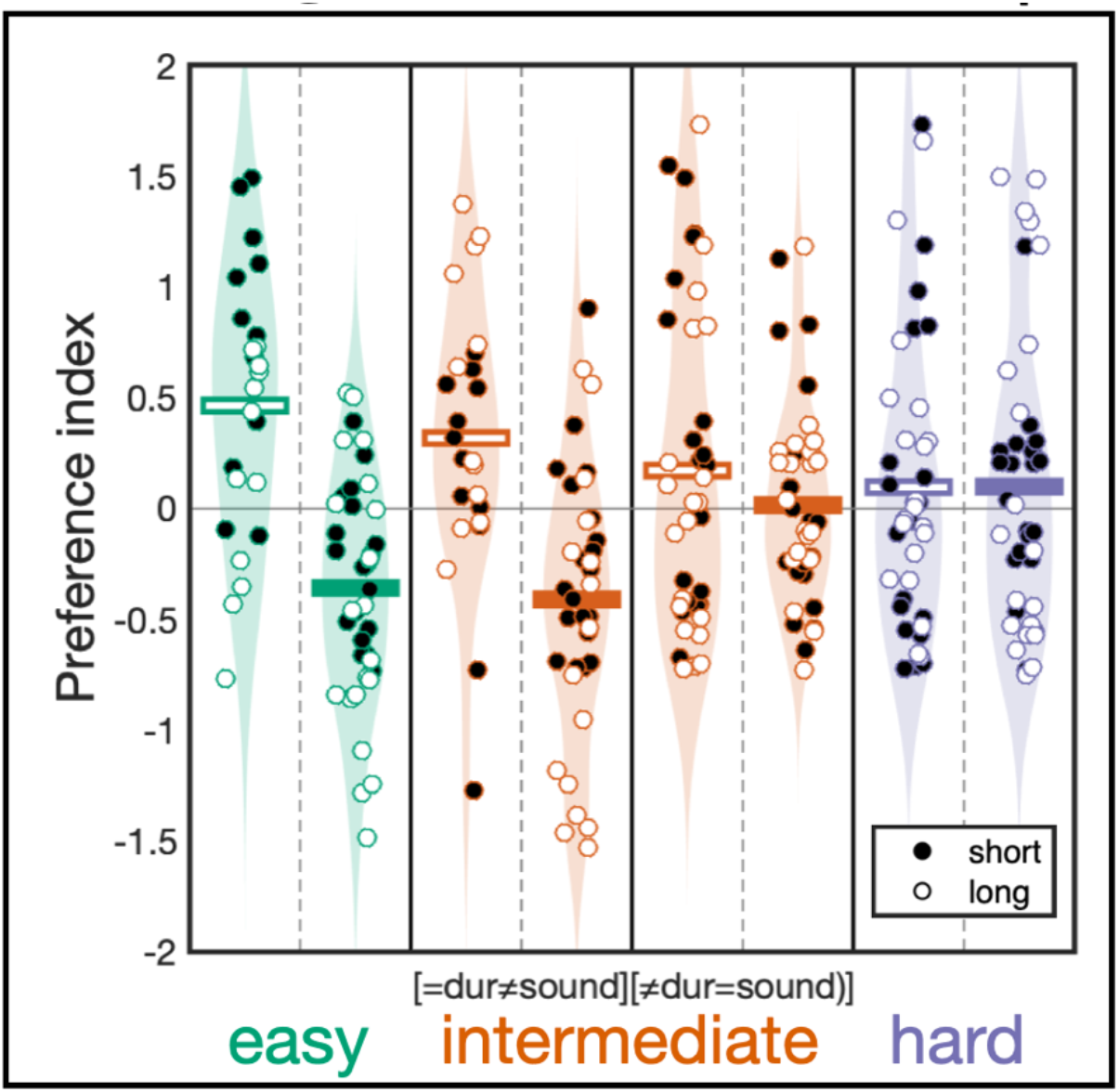
Individual preference in tests. Preference index (mean per subject, n = 284). Filled and empty symbols distinguish imprinting exposure conditions where short or long duration sound assemblages were used, respectively. Coloured patches are kernel density estimations for each vertical column. Horizontal bars represent means for tests corresponding to imprinting exposures to *natural* (open bars) or *artificial* (filled bars) sound assemblages, and are the same data shown in **Figure 2**.

**Extended figure 4.**
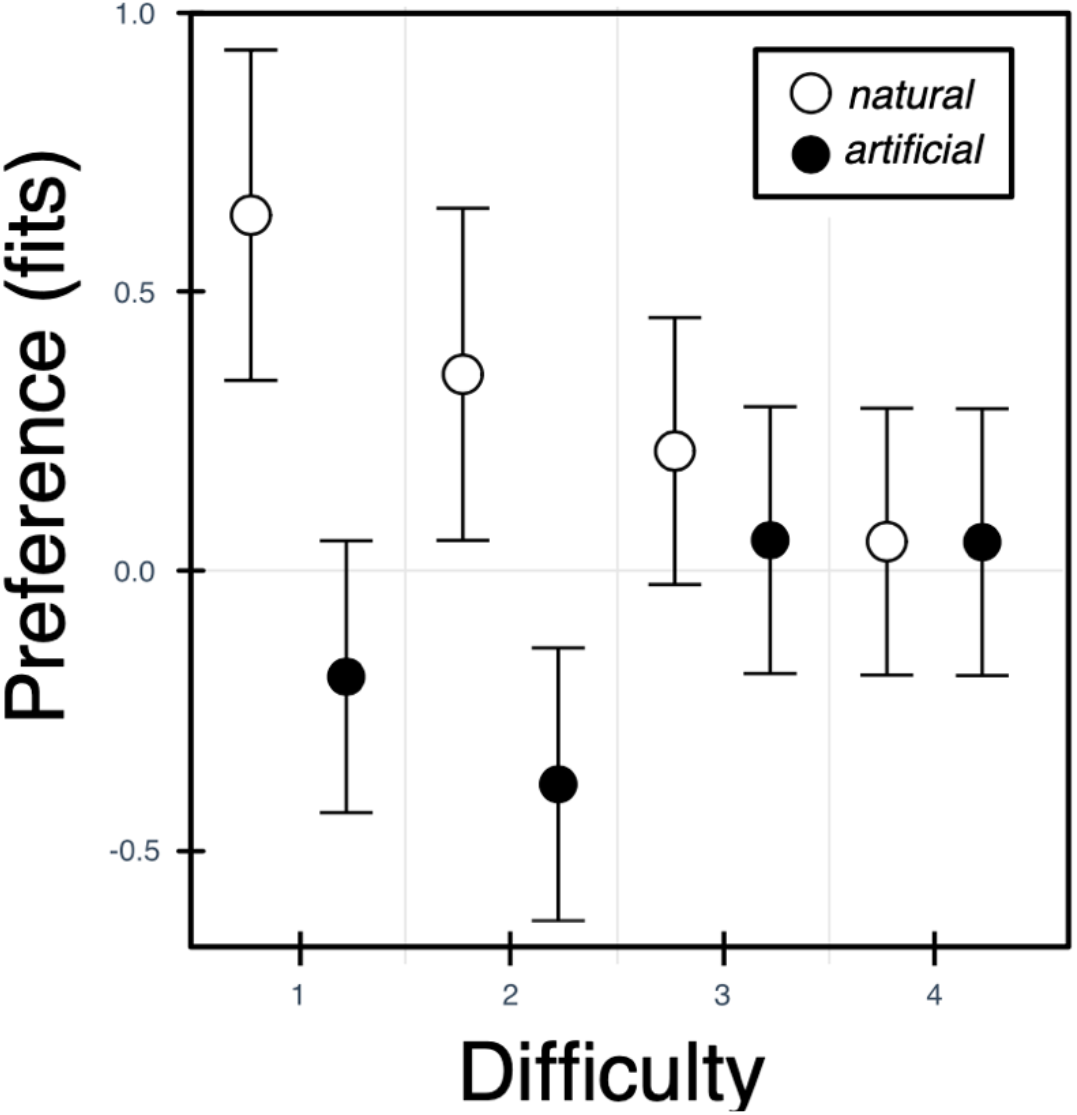
Interaction plot. Fitted values of the preference index (±95% confidence intervals) as a function of test difficulty and sound type used in imprinting exposure.

**Video 1.** Example animal trajectory with overlaid automatic tracking. In this video the target (same stimulus as the one used during imprinting exposure) is coloured in red, competitor in green. This video corresponds to the example animal trajectory depicted in the top left panel of Extended Figure 2.

